# *Packed Like Sardines* – How Surface Crowdedness Impacts Accessibility to Peptidoglycan of *Staphylococcus aureus*

**DOI:** 10.1101/2020.11.10.374892

**Authors:** Noel J. Ferraro, Marcos M. Pires

## Abstract

Bacterial cell walls are essential barriers that protect bacteria against the onslaught of potentially lethal molecules from the outside. Small molecule therapeutics, proteins from bacterial foes, and host immune proteins must navigate past a dense layer of bacterial biomacromolecules (e.g., capsular proteins, teichoic acids, and anchored proteins) to reach the peptidoglycan (PG) layer of Gram-positive bacteria. A subclass of molecules (e.g., antibiotics with intracellular targets) must also permeate through the PG (in a molecular sieving manner) to reach the cytoplasmic membrane. In the case of *Staphylococcus aureus* (*S. aureus*), teichoic acids are the major biopolymers that decorate bacterial cell surfaces. Despite the biological and therapeutic importance of surface accessibility, systematic analyses in live bacterial cells have been lacking. We describe a novel live cell fluorescence assay that reports on the permeability of molecules to and within the PG scaffold. The assay has robust reproducibility, is readily adoptable to any Gram-positive organism, and is compatible with high-throughput sample processing. Analysis of the factors controlling permeability to *S. aureus* and the methicillin resistant MRSA revealed that molecular flexibility plays a central role in molecular permeability. Moreover, teichoic acids impeded permeability of molecules of a wide range of sizes and chemical composition.

## Introduction

Bacterial resistance to antibiotics has become an imminent threat to global health and must be met with an antibiotic pipeline revitalization or, *in lieu* of that, alternative methods to combat bacterial infections.^5,6^ It is well appreciated that lack of membrane permeability can negatively impact the antibacterial activity of small molecules with intracellular (or periplasmic) targets.^1–3^ The role of accessibility to therapeutic targets on bacterial cell surfaces – that do not require membrane penetration – remains far less defined and has not been systematically characterized. The usual presumption is that molecular targets on extracellular regions of bacteria are readily accessible to antibacterial molecules and proteins. However, due to the dense brush-like macromolecules (teichoic acids) that decorate the surface and the matrix-like peptidoglycan (PG) layer, therapeutic targets on bacterial cell surfaces could potentially be difficult to reach. (**Figure 1**). The therapeutic effectiveness of vancomycin (and a multitude of other antibiotics with the same target) hinges on its ability to permeate through bacterial biomacromolecules to ultimately reach lipid-anchored target. However, bacteria can leverage this feature of impeded accessibility to escape the action of antibacterial agents like vancomycin. For example, *Staphylococcus aureus* (*S. aureus*) resistance to vancomycin can result from cell wall thickening, which effectively captures vancomycin and prevents its association with lipid II.^4^ A molecular sieving concept through the dense cell wall has also been evoked to describe trends in antibacterial activities of synthetic mimics of antimicrobial peptides.^5^ Similarly, essential components of the human innate and adaptive immune system such as lysozyme and antibodies, respectively, target cell surface components and do not need to cross the membrane bilayer but have been met with resistance potentially due to effects concerning accessibility.

**Figure 1.**
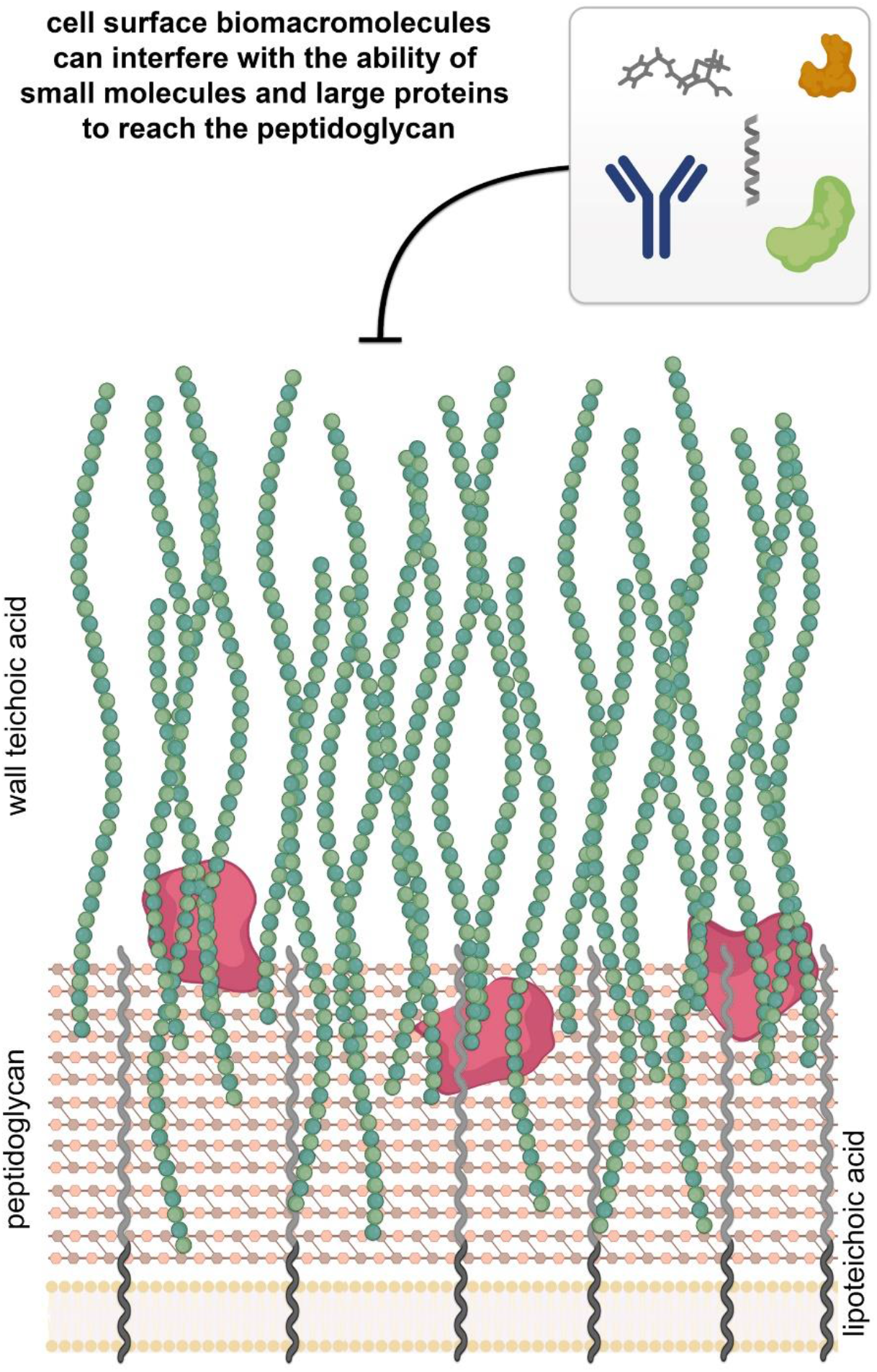
Surface composition can impact accessibility of molecules to the PG scaffold. Schematic representation of the surface composition of *S. aureus* delineating key biomacromolecules that can potentially impact penetration of molecules.

The composition of surface-bound macromolecules varies across species and has evolved to closely match the ecological niche of that bacterial species. Even in the way bacteria are grouped as Gram-positive, Gram-negative, and mycobacteria has a principle link to their surface architecture. Gram-positive bacteria have a cell wall that includes a thick PG layer on the exterior side of the cytoplasmic membrane (**Figure 1**). PG is a mesh-like polymer made up of repeating disaccharides *N*-acetylglucosamine (GlcNAc) and *N*-acetylmuramic acid (MurNAc). Each MurNAc unit is connected to a short and unusual peptide (stem peptide) with the canonical sequence of L-Ala-iso-D-Glu-L-Lys-D-Ala-D-Ala or *meso*-diaminopimelic acid (*m*-DAP) in place of L-Lys at the 3^rd^ position.^6^ Bacterial PG is an essential component of the bacterial cell wall, which makes it an attractive target for the innate immune system and a large number of FDA-approved antibiotics.^7, 8^ For all known bacteria, neighboring stem peptides are crosslinked to endow the PG matrix with rigidity and integrity. Cell walls are further decorated with a number of polymers and proteins that play important physiological roles such as regulating cell morphology and growth, serving as virulence factors, and aiding in adherence and colonization (**Figure 1A**).^9^

Among Gram-positive pathogens, *S. aureus* has proven to be particularly challenging to combat due to its various mechanisms of antibiotic resistance and host immune evasion. *S. aureus* is most commonly observed in skin infections but it can cause more severe internal infections.^11,12^ According to the Centers for Disease Control and Prevention (CDC), hard-to-treat methicillin resistant strains of *S. aureus* (MRSA) infect an estimated 120,000 people and cause ∼20,000 deaths a year in the United States alone. The difficulty in finding new efficacious antibiotics against *S. aureus* highlights the need to explore less conventional therapeutic approaches such as antibiotic adjuvants or immunotherapies.^10–15^ For example, adjuvants can potentiate antibiotics by improving their accessibility to their cognate molecular target. Likewise, anti-infective immunotherapeutics (e.g., antibody recruiting agents developed by our lab^10–13, 15^) work by targeting macromolecules on bacterial cell surfaces. Despite the pivotal role of surface accessibility in bacterial pathogenesis, to date there has not been a systematic characterization of how molecular size and flexibility control penetration of biomacromolecules across bacterial cell surfaces; information that can directly impact the improvement of current therapies and the development of alternative treatments. We describe a facile assay that reports on surface features that most critically impact the access of molecules to the PG scaffold of live *S. aureus* (and is potentially adaptable to any other Gram-positive bacterium).

## Results and Discussion

Early seminal works have described a molecular sieving effect of polymers permeating through bacterial PG, which is likely a product of its lattice structure.^16, 17^ While illuminating, these experiments were performed *in vitro* with isolated PG (sacculi). We set out to develop a method to systematically measure accessibility to the PG scaffold of live bacterial cells. The basis of the assay is a site selective incorporation of a reactive epitope within the PG of live cells followed by treatment with heterobifunctional reporter molecules of varying sizes that attach to the PG scaffold (**Figure 2A**). The reporter molecules are linked to fluorophores, and, therefore, cellular fluorescence levels describe the ability of the probes to navigate through surface exposed biomacromolecules. Covalent PG tagging should result in reliable measurements that can be readily quantified using standard techniques amendable to high throughput analyses (e.g., flow cytometry). We initially reasoned that a thiol functional group could be installed within bacterial PG, which lacks native thiols, by the process of metabolic tagging of the PG with synthetic stem peptide analogs containing D-cysteine.^11, 18–21^ During cell growth and division, synthetic PG analogs enter the biosynthetic pathway in place of endogenous building blocks, and this process provides a robust route to introduce non-native functional groups within bacterial PG.^22^

**Figure 2.**
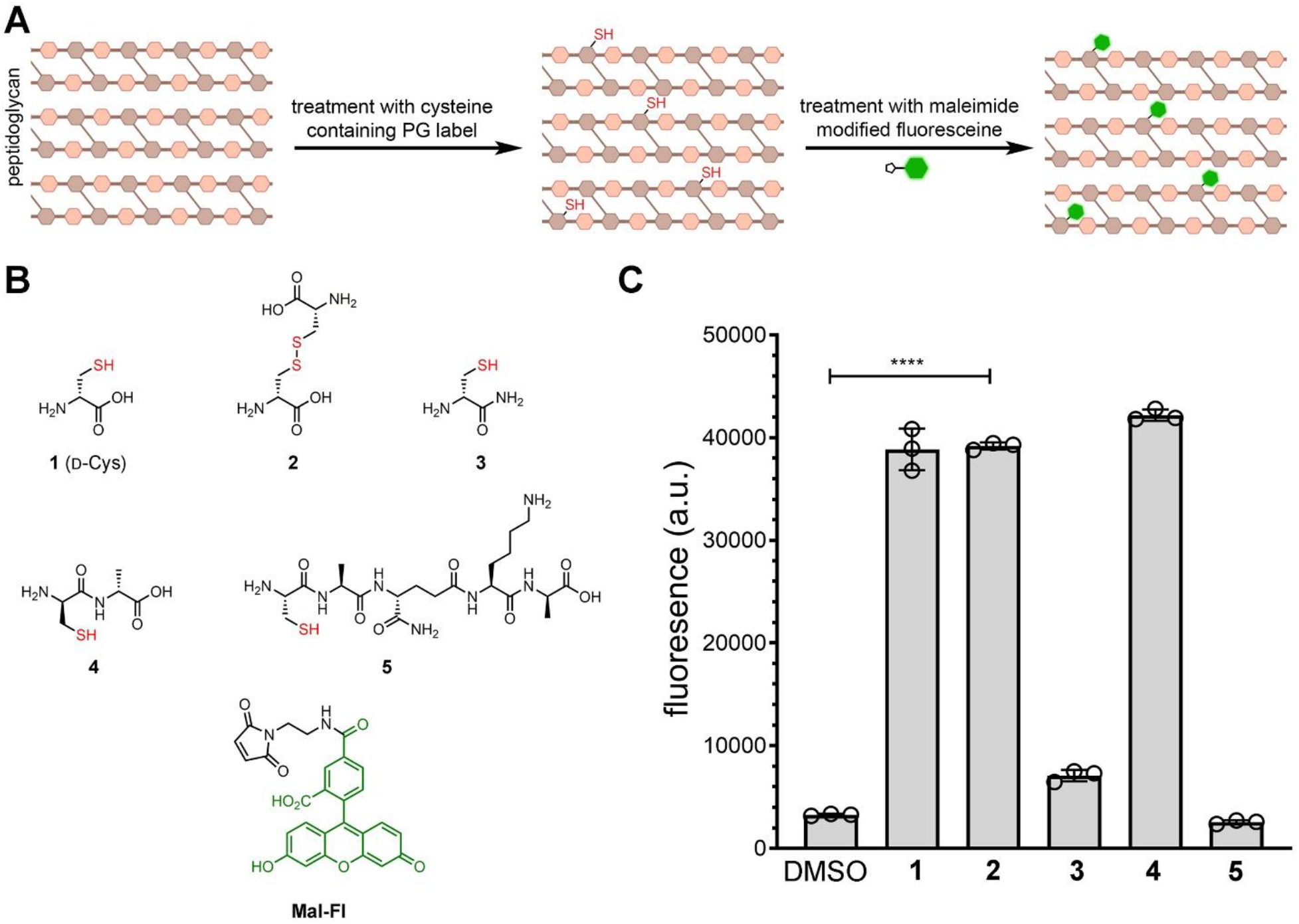
Bench-marking assays to show accessibility to bacterial PG scaffold. **(A)** Proposed assay to tag the bacterial PG scaffold with thiol handles followed by a fluorescent probe that contains an orthogonal binding partner for the thiol handle. **(B)** Chemical structure of PG analogs modified with a cysteine residue and the fluorescent reporter **Mal-Fl. (C)** Flow cytometry analysis of *S. aureus* (ATCC 25923) treated overnight with 1 mM of synthetic PG analogs, reduced with DTT (5 mM), and incubated with 25 μM of **Mal-Fl**. Data are represented as mean +/- SD (n = 3). *P*-values were determined by a two-tailed *t*-test (* denotes a *p*-value < 0.05, ** < 0.01, ***<0.001, ns = not significant).

A small panel of synthetic PG analogs was synthesized, each of which contained a cysteine residue (**Figure 2B**). The panel contained three derivatives of the single D-amino acid, D-cysteine. During cell growth, exogenously supplied single D-amino acids, such as D-cysteine, are swapped in the place of the D-alanine that occupies the 5^th^ position within the stem peptide of *S. aureus* (**Figure S1**). A wide range of single D-amino acid PG probes have been developed to elucidate fundamental steps in bacterial cell wall biology.^13, 22–32^ We also included oxidized D-cystine (**2**) and D-cysteine amidated in the *C*-terminus (**3**) in an attempt to maximize PG tagging.^26, 30^ Cell treatment with the dipeptide D-Cys-D-Ala (**4**) was expected to result in the incorporation of D-cysteine at the 4^th^ position of the stem peptide.^11, 33, 34^ D-Cys-D-Ala mimics the D-Ala-D-Ala dipeptide PG precursor, therefore it should be processed intracellularly by the MurF enzyme to generate D-cysteine containing PG. Lastly, cysteine was placed at the *N*-terminus of a tetrapeptide synthetic analog (**5**) of the PG stem peptide. We^32, 35^, and others^36–39^, recently showed that structural analogs of PG stem peptides can be crosslinked into the growing PG scaffold of live cells.

Our first goal was to identify which cysteine-based label would result in the highest level of thiol handles on the surface of *S. aureus*. To accomplish this, *S. aureus* cells were grown overnight in the presence of each PG analog to promote incorporation throughout the entire PG scaffold. Then, cells were treated with the reducing agent dithiothreitol (DTT) to unmask the thiols on the PG which were expected to exist primarily as disulfides due to the oxidizing culture media. Cells were washed with PBS to remove excess reducing agent and incubated with maleimide-modified fluorescein (**Mal-Fl**). Our results clearly showed that some of the PG metabolic tags resulted in significant increases in fluorescence levels (**Figure 2C**). Incubation of cells with D-cystine (**2**) led to a ∼12-fold increase in cellular fluorescence relative to DMSO treated cells. Likewise, treatment of *S. aureus* cells with the enantiomeric L-cystine, which is not expected to be processed by PG transpeptidases, led to a minimal increase in cellular fluorescence compared to DMSO treated cells (**Figure S2**). Next, a titration experiment was carried out and we found that 25 μM was an optimum concentration of **Mal-Fl** (**Figure S3**). Although there was similar labeling levels observed with the other PG analogues, we expected that D-cystine, the disulfide form of D-cysteine, would result in more consistent labeling levels compared to the other compounds due to the fact that the others may exist at varying levels of oxidized product during the incubation period in the oxygenated media. Based on these results, we selected the PG probe D-cystine for all subsequent assays that use the thiol-maleimide pair.

Localization studies were performed next to show that the **Mal-FI** is imbedded within the bacterial PG scaffold after reacting with the thiol handle. Our laboratory had previously demonstrated that treatment of *S. aureus* with single D-amino acid probes resulted in the chemical modification of the stem peptide using a range of biochemical techniques.^11, 28, 29^ Nonetheless, *S. aureus* cells were treated with D-cystine followed by **Mal-Fl** and visualized by confocal microscopy. Our results showed that the reporter probe had similar labeling profile as *S. aureus* labeled with the single amino acid probe **D-LysFl** (**Figure 3A**). The same cells were subjected to a sacculi isolation procedure, which were similarly imaged by confocal microscopy. Satisfyingly, the pattern of labeling in the isolated PG scaffold mirrored that of the whole cell and are suggestive of the probe being covalently attached to the PG scaffold. A second line of evidence of PG binding by the accessibility probe was provided by treatment of *S. aureus* with digestion enzymes (**Figure 3B**). As before, *S. aureus* were labeled with D-cystine followed by **Mal-Fl** and chemically fixed to preserved the overall cellular structures for flow cytometry analysis. Fixed cells were subjected to treatment with either proteinase K or mutanolysin and cellular fluorescence was measured periodically. Mutanolysin is a muralytic enzyme that cleaves the polysaccharide backbone of PG, thus triggering the release of PG fragments from the cell. Fluorescent probes that are covalently attached to the PG should, likewise, separate from the cells upon treatment with mutanolysin, leading to a reduction in cellular fluorescence levels. As expected, *S. aureus* cells treated with mutanolysin demonstrated a fluorescent level that was half that of the starting level by 120 minutes. Conversely, fluorescence levels of *S. aureus* cells treated with proteinase K (a promiscuous protease that typically cleaves the peptide bond adjacent to aliphatic and aromatic amino acids) remained mostly unchanged over the course the entire experiment. Together, these results are strongly suggestive of the modification of bacterial PG following the two-step labeling procedure (installation of D-cystine within the PG scaffold followed by covalent attachment of the accessibility probe).

**Figure 3.**
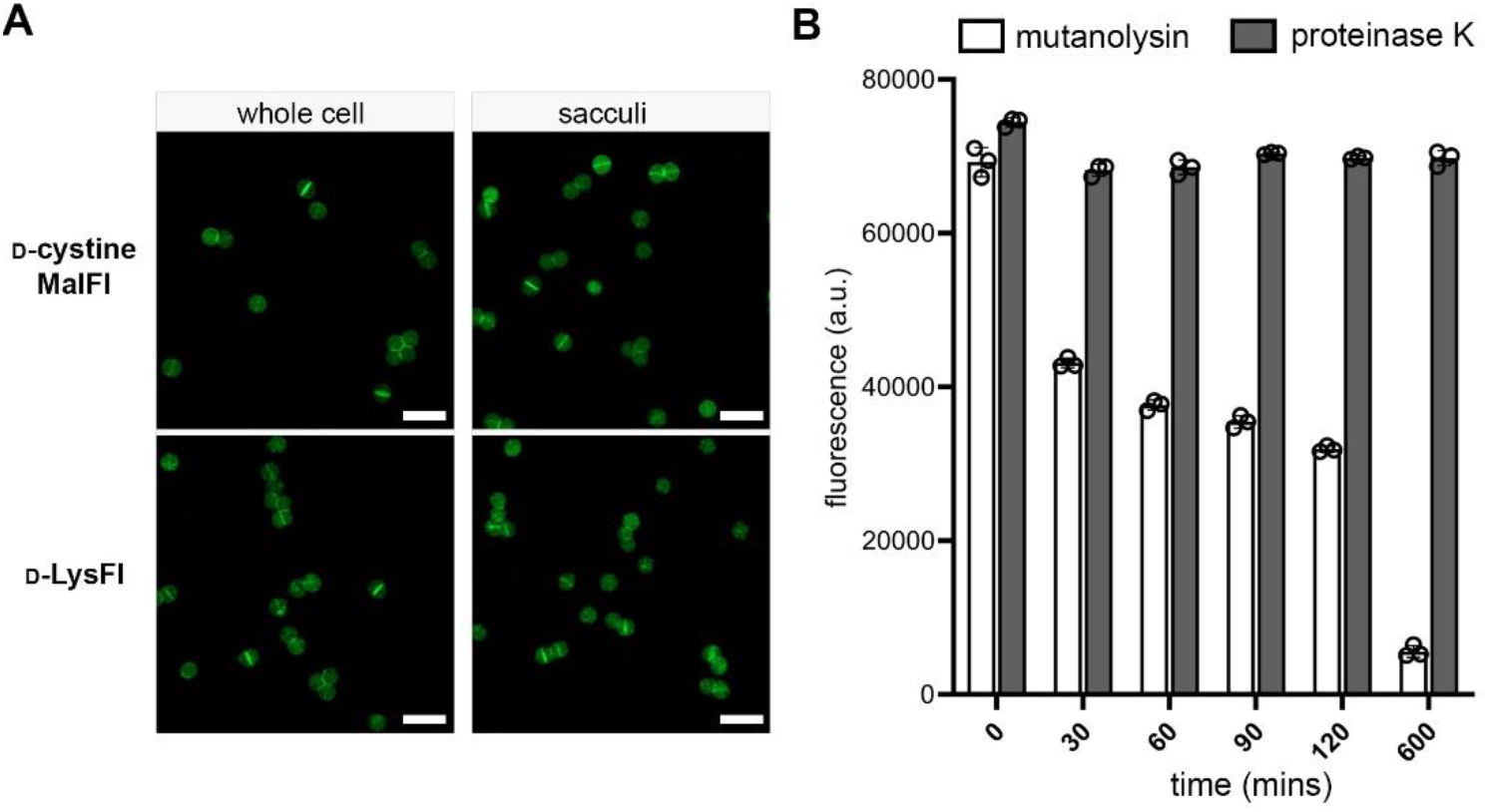
**(A)** Confocal microscopy of *S. aureus* or isolated sacculi from *S. aureus. S. aureus* (ATCC 25923) treated overnight with 1 mM of D-cystine or **D-LysFl**. Cells treated with D-cystine were reduced with DTT (5 mM) and incubated with 25 μM of **Mal-Fl** and imaged. Scale bar = 5 mm. Cells were either imaged by confocal microscopy or subjected to a secondary step in which the sacculi was isolated and imaged. **(B)** Flow cytometry analysis of surface labeled *S. aureus* (ATCC 25923) treated proteinase K (dark bars) or mutanolysin (clear bars). As in (A), surface labeled cells were treated with 1 mM of D-cystine, reduced with DTT (5 mM), and incubated with 25 μM of **Mal-Fl**. Data are represented as mean +/- SD (n = 3). *P*-values were determined by a two-tailed *t*-test (* denotes a *p*-value < 0.05, ** < 0.01, ***<0.001, ns = not significant).

Accessibility to the bacterial PG layer and permeation within the PG scaffold by molecules from the extracellular space should be tied to their physiochemical properties (e.g., charge, size, and flexibility). To test these concepts, we assembled two libraries of accessibility probes that, like **Mal-Fl**, display maleimide and fluorescein functional groups. One library contained a flexible polar polyethylene glycol (PEG) spacer and the other a rigid polyproline spacer of varying lengths (**Figure 4**). As before, the PG scaffold of *S. aureus* cells was tagged with thiol handles by incubating with D-cystine. Accessibility to the PG scaffold was investigated by treating cells with members of both libraries and was analyzed by flow cytometry. A stark difference in surface accessibility was noted between the two libraries. Increasing the length of the spacer in **Mal-peg**_**n**_**-Fl** resulted in a gradual and consistent decrease in fluorescence, whereas there was a sharp decrease in cellular fluorescence with the series of **Mal-pro**_**n**_**-Fl**. These results indicate that rigidity of a molecule may not be favorable for reaching the PG scaffold of bacteria. Instead, flexibility may promote the maneuvering of molecules across surface biopolymers. A similar labeling profile was observed with two other strains of *S. aureus*, including the MRSA strain USA300 (**Figure S4**). These results likely reflect the conserved nature of D-cystine labeling in these organisms and could make this assay applicable to other *S. aureus* strains.

**Figure 4.**
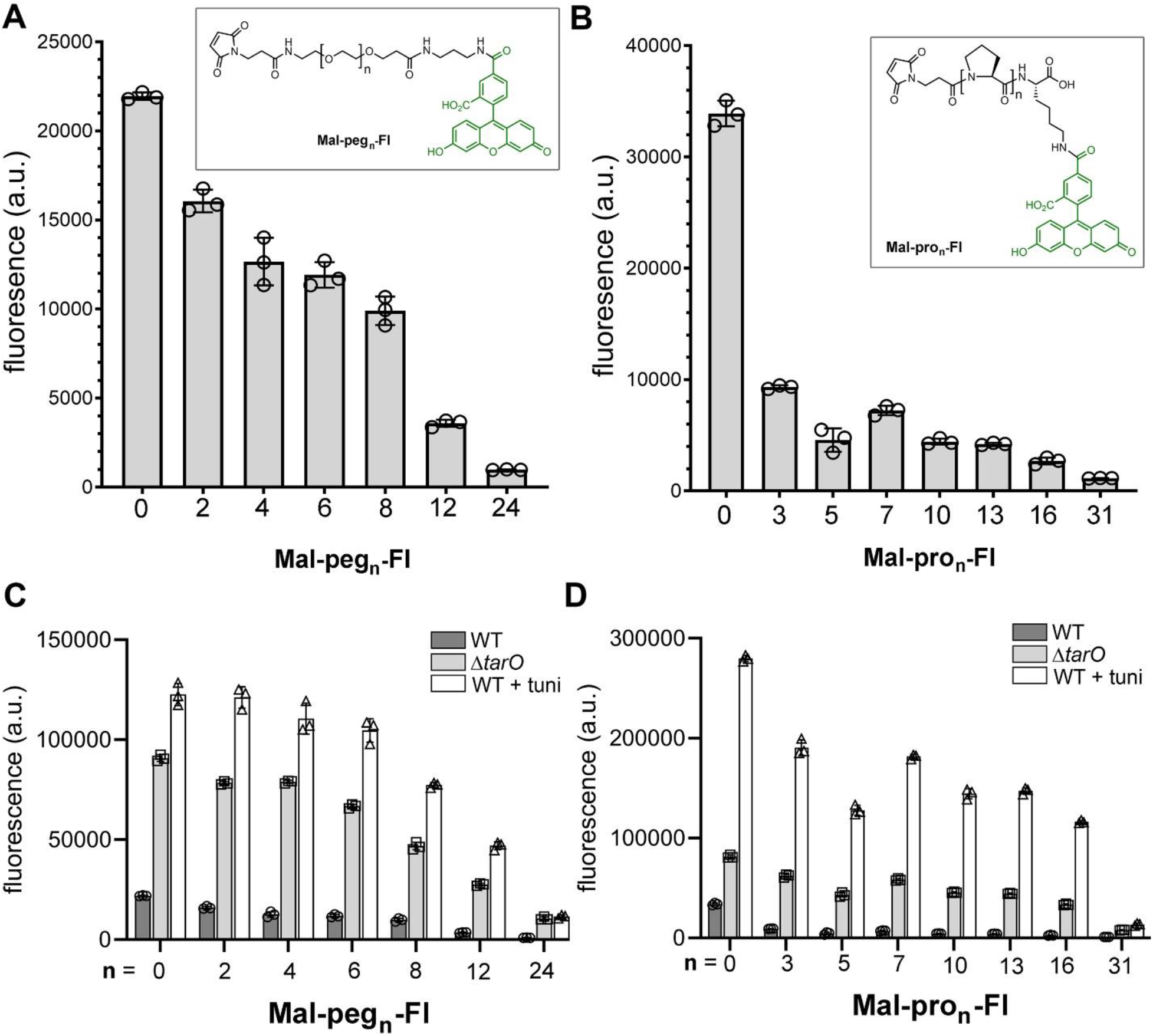
**(A-B)** Flow cytometry analysis of WT *S. aureus* (ATCC 25923) treated overnight with 1 mM of D-cysteine, reduced with DTT (5 mM), and incubated with 25 μM of designated accessibility probes. (**C-D**) WT *S. aureus* (ATCC 25923) cells were incubated with 1 mM of D-cystine (dark bars), co-incubated with tunicamycin (tuni, 0.1 μg/mL) and 1 mM of D-cystine (grey bars), or *S. aureus* (*ΔtarO*) were incubated with 1 mM of D-cystine alone (white bars) overnight. Next, cells were reduced with DTT (5 mM), and incubated with 25 μM of designated accessibility probes. Data are represented as mean +/- SD (n = 3).

We proceeded to investigate the role of surface biopolymers on PG accessibility. There are two main surface biopolymers on *S. aureus* cells known as lipoteichoic acids (LTA) and wall teichoic acids (WTA).^40–43^ WTA is highly anionic and forms a dense glycan layer that is covalently attached to the stem peptide (**Figure 1**). Previous reports have described the influence of WTA on bacteriophage susceptibility,^44^ antibody binding,^45^ antibiotic resistance (e.g., daptomycin),^46^ and recognition by innate immune proteins that bind to PG (e.g., Peptidoglycan Recognition Protein).^47^ Inhibitors of WTA biosynthesis, such as tunicamycin, have been developed as anti-infective agents.^48–51^ Tunicamycin inhibits TarO, which is responsible for the first step in WTA biosynthesis.^52^ We sought to gain a more systematic description of the impact of WTA on accessibility to *S. aureus* PG by using a *tarO* deletion strain and tunicamycin-based WTA inhibition. Both modes of WTA disruption resulted in large increases in cellular labeling with the accessibility probes (**Figure 4 C-D)**. For example, fluorescence levels in *S. aureus* (*tarO*) treated with the 36-atom long spacer, **Mal-peg**_**12**_**-Fl**, was higher than *S. aureus* (WT) with no spacer. Tunicamycin treatment yielded even higher levels of cellular fluorescence. The boost in cellular fluorescence was more pronounced with rigid spacers as demonstrated by the ratio of cellular fluorescence when treated with **Mal-pro**_**13**_**-Fl** and **Mal-Fl**. In the absence of tunicamycin treatment, cellular fluorescence of **Mal-pro**_**13**_**-Fl** was ∼6.5% relative to **Mal-Fl** and this ratio jumped to ∼40% upon tunicamycin treatment. These results are consistent with WTA playing a determinant role is preventing rigid molecules from reaching the bacterial cell surface.

We next set out to test how robust the concept of this assay is by changing the reactive partners. Instead of thiol and maleimide, we assembled a panel of probes centered on an azide-modified D-lysine (**D-LysAz**) and DiBenzoCycloOctyne (DBCO) conjugated to a fluorescent handle (**Figure 5A**).^53, 54^ This pair of reactive functional groups is biorthogonal and readily forms a triazole covalent bond in the absence of metal catalysts. We found that *S. aureus* cells incubated with **D-LysAz** followed by treatment with DBCO-Fl resulted in a ∼12-fold increase in cellular fluorescence compared cells not treated with the unnatural D-amino acid (**Figure 5B**). Moreover, the fold increase in cellular fluorescence was very similar to the values observed using the D-cystine and **FITC-Fl** pair. Cellular treatment with the enantiomer **L-LysAz** led to background fluorescence levels, another indication that fluorescence signals represent covalent modification of the PG scaffold (**Figure S6**). In addition, mutanolysin analysis revealed a similar profile (**Figure S7**) and confocal microscopy showed identical localization pattern (**Figure S8**) to the thiol-maleimide pair.

**Figure 5.**
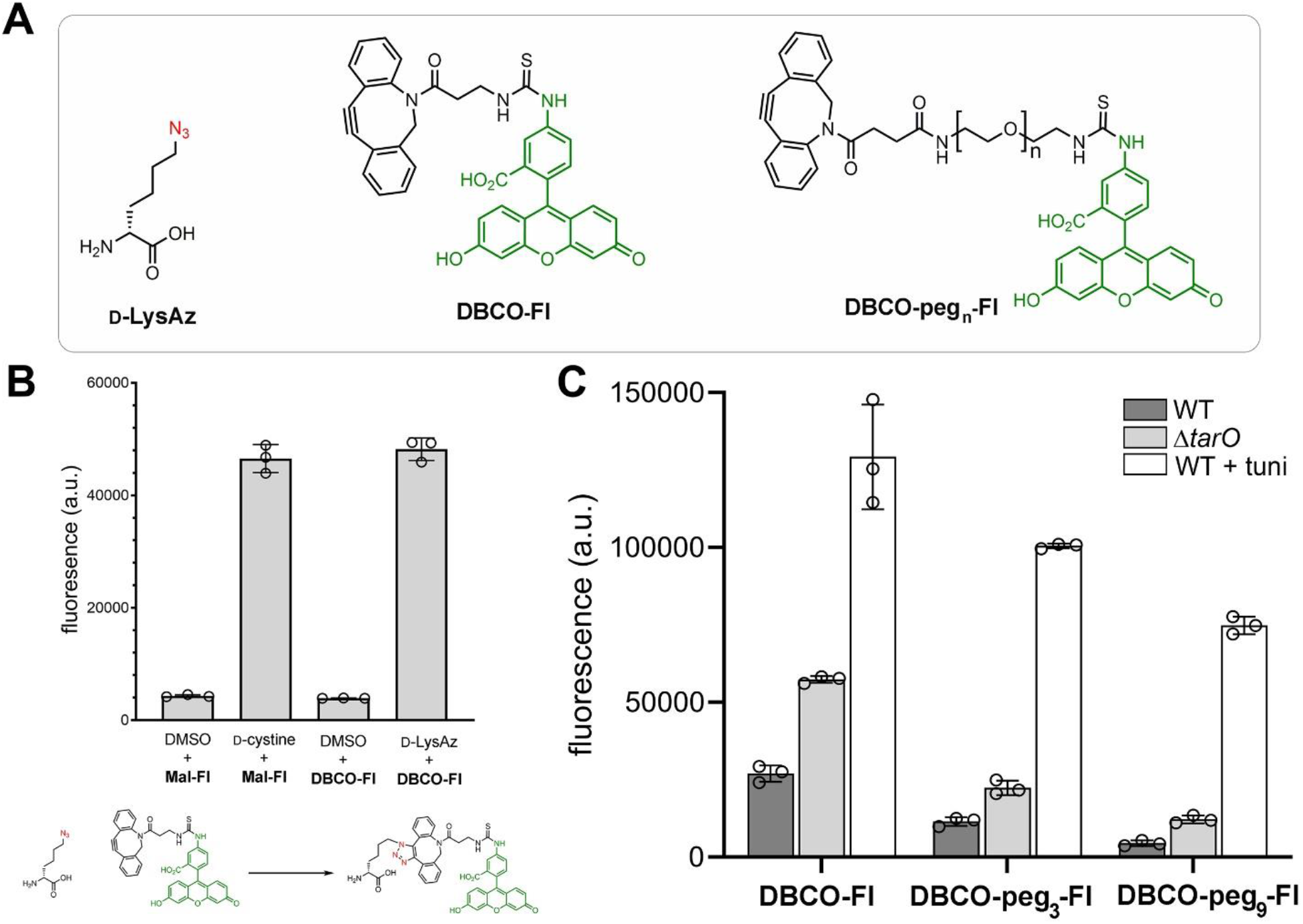
**(A)** Chemical structures of **D-LysAz, DBCO-Fl**, and **DBCO-peg**_**n**_**-Fl. (B)** Flow cytometry analysis of WT *S. aureus* (ATCC 25923) treated overnight with 1 mM of D-cystine or **D-LysAz** followed by a treatment with either 25 μM of **Mal-Fl** or **DBCO-Fl. (C)** WT *S. aureus* (ATCC 25923) cells were incubated with 1 mM of **D-LysAz** alone (dark bars), co-incubated with tunicamycin (tuni, 0.1 μg/mL) and 1 mM of **D-LysAz** (grey bars), or *S. aureus* (*ΔtarO*) were incubated with 1 mM of **D-LysAz** alone (white bars) overnight. Next, cells were treated with 25 μM of designated accessibility probes. Data are represented as mean +/- SD (n = 3).

The effect of WTA on PG accessibility was evaluated for the biorthogonal pair (**Figure 5C**). Loss of WTA in the deletion strain resulted in a two-fold increase in cellular fluorescence across all spacer lengths. Co-incubation of cells with tunicamycin resulted in ∼9-fold increase in cellular fluorescence with the **DBCO-peg**_**3**_**-Fl** probe and a ∼17-fold increase with the longest **DBCO-peg**_**9**_**-Fl** probe. These results suggest that the assay platform is robust and the chemistry is not a factor in measuring access to the PG scaffold. Next, we sought to evaluate the role of branched polyethylenimine (BPEI) in potentiating β-lactam antibiotics against MRSA.^55, 56^ Whereas *tarO*-null strain and tunicamycin pre-treatment results in cells lacking WTA, BPEI has been proposed to directly interact with WTA without inhibiting its biosynthesis. Binding of BPEI to WTA was suggested to result in the synergy of β-lactam antibiotics by causing delocalization of penicillin binding proteins. We reasoned that neutralization of WTA by BPEI could also alter accessibility of molecules to the PG scaffold. To test this, *S. aureus* cells were treated with BPEI and challenged with our probes (**Figure 6A**). Our results reveal that BPEI improved the accessibility of smaller molecules to the PG, which likely plays a role in its synergistic activity with small molecule antibiotics. The increased accessibility appeared to be size dependent, as larger molecules did not permeate better when co-incubated with BPEI. Finally, we set out to assess the effect of LTA on surface accessibility.^57, 58^ Unlike WTA, LTA is anchored into the bacterial membrane *via* a glycolipid group. Although the roles of LTA have not been fully elucidated, LTA has been implicated in a diverse set of functions including interaction with host toll-like receptors,^59^ organization of cell division machinery,^60, 61^ and regulating biofilm formation.^62^ The gene responsible for LTA biosynthesis, *ltaS*, is essential for growth of *S. aureus*^61^ but becomes conditionally essential when the chaperon ClpX is inactivated.^63^ In our assay, we found that accessibility to the PG of cells lacking LTA was significantly increased (**Figure 6B**). Deletion of *clpX* alone did not alter the permeation of **DBCO-Fl**, which indicates that deletion of ClpX alone cannot account for the increased permeability. We then investigated the role of LTA D-alanylation on surface accessibility using a small molecule inhibitor, amsacrine, that was previously described.^49^ Introduction of a positively charged D-alanine within LTA results in the neutralization of the anionic teichoic acids. This modification has been shown to influence a number of biological functions including biofilm formation,^64, 65^ sensitivity to antibiotics (e.g., antimicrobial peptides and daptomycin),^57, 66–68^ and recognition by human innate immune system.^69^ However, its role in surface accessibility has not been evaluated. We found that treatment of *S. aureus* with amsacrine resulted in a modest increase in cellular fluorescence with **DBCO-peg**_**3**_**-Fl** (**Figure S9**). Collectively, these results provide direct evidence that chemical or biochemical alterations to teichoic acids on the surface of bacteria can regulate the permeation of molecules. In turn, these results may reveal a new facet to the diverse ways that bacteria modulate access to essential components of the cell wall.

**Figure 6.**
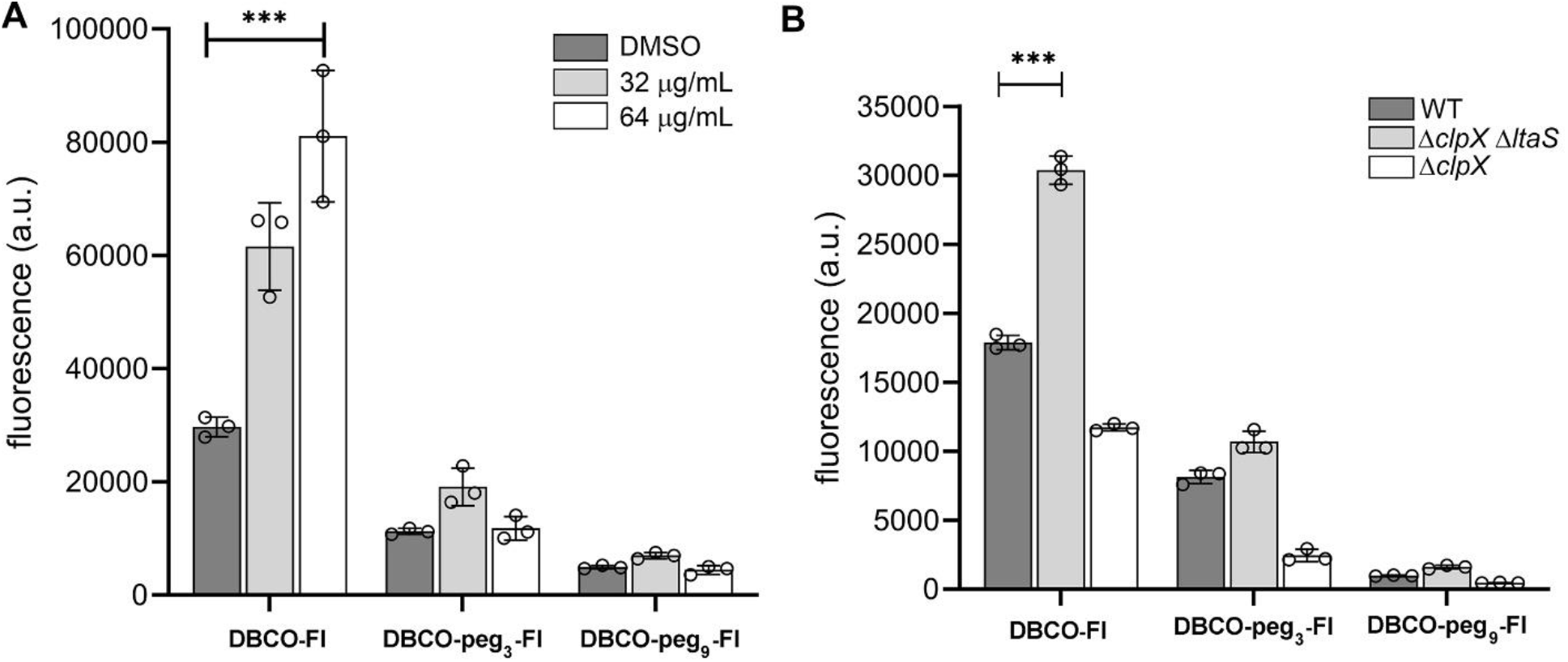
**(A)** Flow cytometry analysis of WT *S. aureus* (ATCC 25923) treated overnight with 1 mM of **D-LysAz**, co-incubated with 0, 32, or 64 μg/mL of BPEI at stationary phase, followed by a treatment with 25 μM of designated probes. **(B)** Flow cytometry analysis of WT *S. aureus* (ATCC 25923), *S. aureus* (*ΔclpX ΔltaS*), or *S. aureus* (*ΔclpX*) treated overnight with 1 mM of **D-LysAz**, followed by a treatment with 25 μM of designated probes. Data are represented as mean +/- SD (n = 3). *P*-values were determined by a two-tailed *t*-test (* denotes a *p*-value < 0.05, ** < 0.01, ***<0.001, ns = not significant).

## Conclusion

In conclusion, we have developed a novel fluorescence-based assay that reports on the accessibility of molecules to the surface of bacteria. Using *S. aureus* as a model organism, we showed that two distinct chemical handles (thiol and azide) were installed within the PG scaffold of *S. aureus*. Using two focused libraries in which each member contained a reactive handle and a fluorophore, we were able to show the effect of molecular size and flexibility on cellular accessibility. Molecules that are rigid, such as polyproline, displayed low access to the bacterial cell surface. Moreover, the presence of WTA (and to a less extent LTA), played a central role in regulating surface accessibility. Together, these results demonstrate that the assay outlined here is robust, potentially widely adaptable, and can play a significant role in elucidating dynamic features of bacterial cell surfaces.

