## Supplemental Figures for "*Packed Like Sardines* – How Surface Crowdedness Impacts Accessibility to Peptidoglycan of *Staphylococcus aureus*"

### Supporting Figures

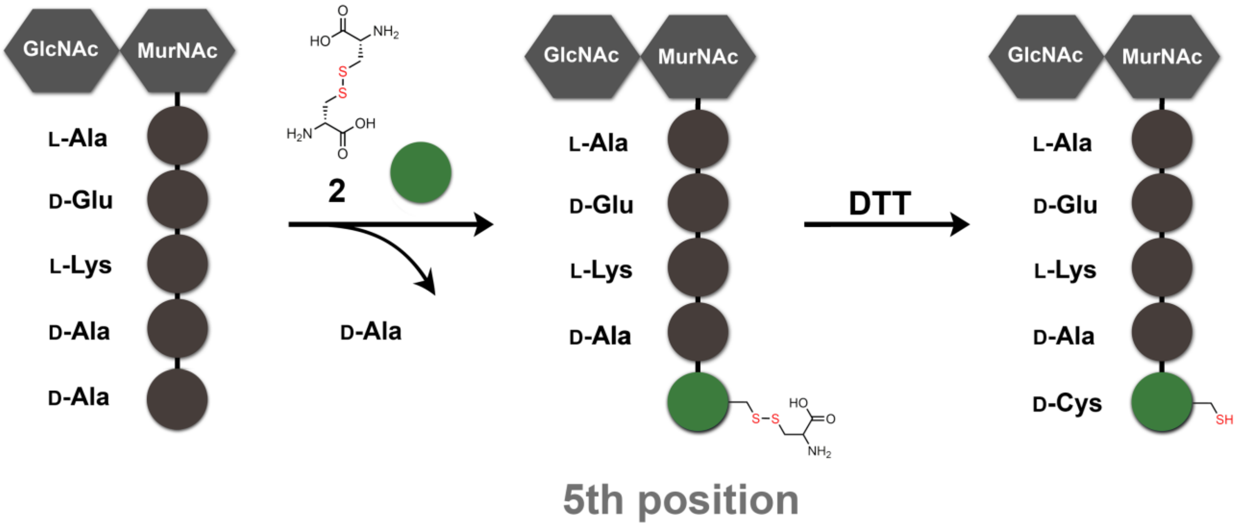


**Figure S1.** Modes of incorporation of single amino acids by swapping into the 5th position on the peptidoglycan stem peptide of *S. aureus* cells. D-cystine is incorporated (singly or doubly) within the PG scaffold and is subsequently reduced with DTT to generate D-cysteine.

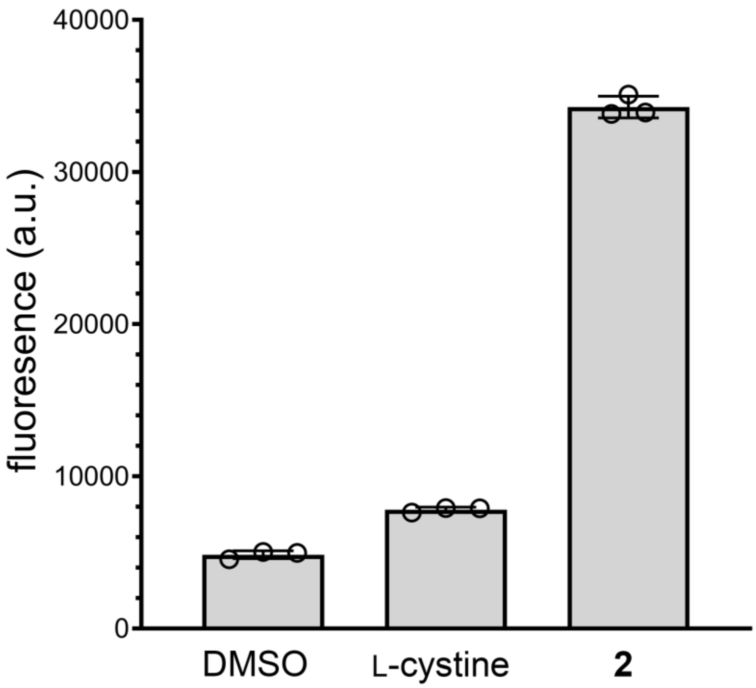


**Figure S2.** Flow cytometry analysis of *S. aureus* (ATCC 25923) treated overnight with 1 mM of DMSO/L-cystine/D-cystine, reduced with DTT (5 mM), and incubated with 25 μM of **Mal-Fl**. Data are represented as mean +/- SD (n = 3).

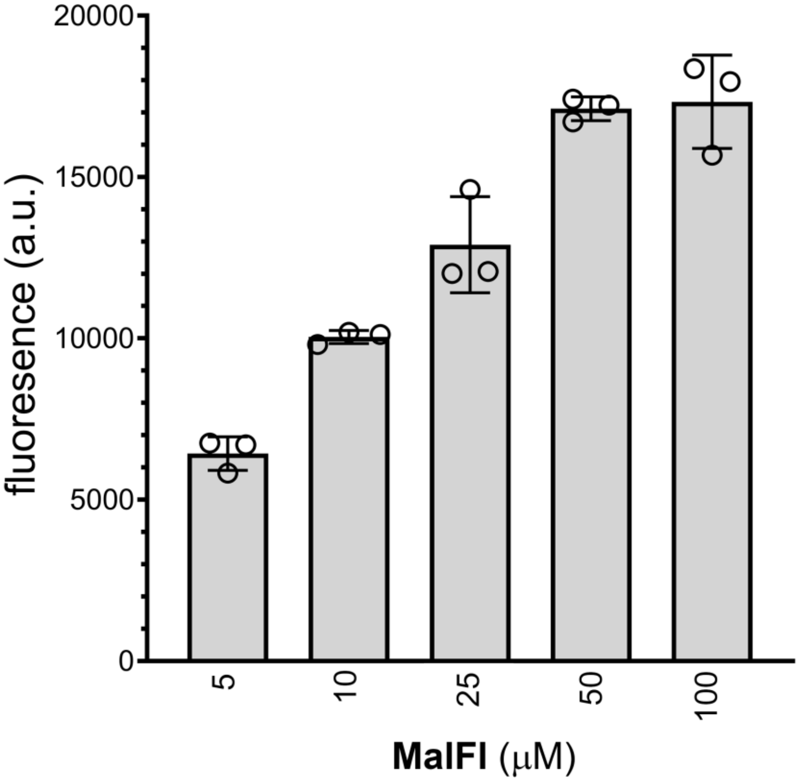


**Figure S3.** Flow cytometry analysis of *S. aureus* (ATCC 25923) treated overnight with 1 mM of D-cystine, reduced with DTT (5 mM), and incubated with increasing concentrations **Mal-Fl**. Data are represented as mean +/- SD (n = 3).

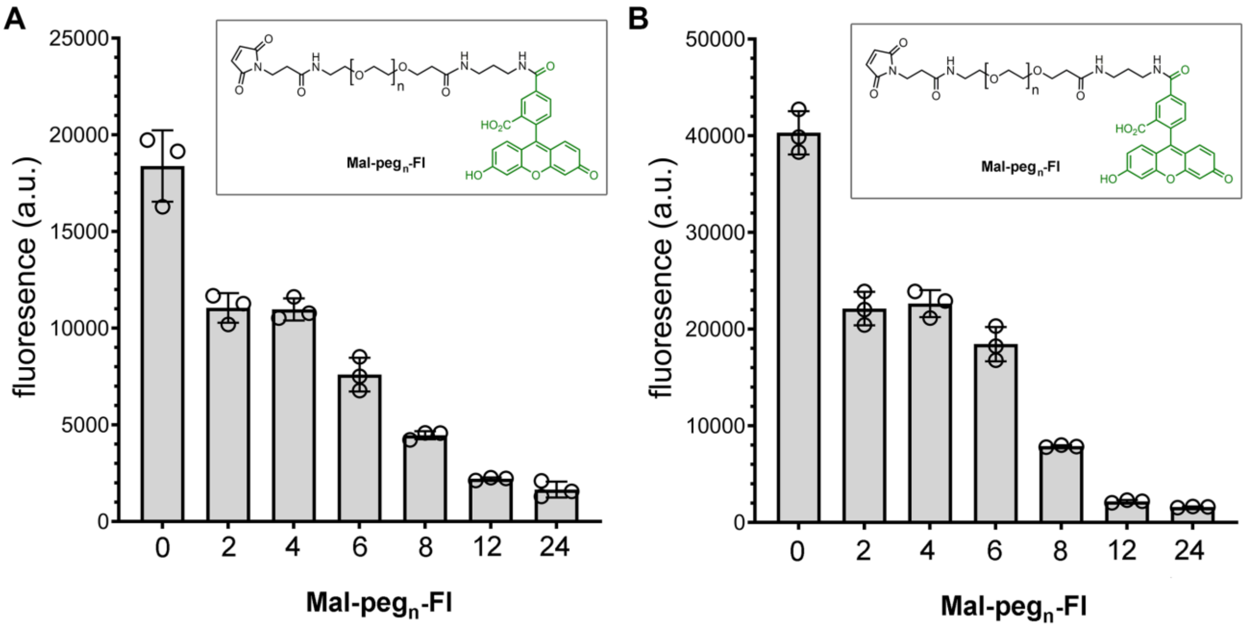


**Figure S4.** Flow cytometry analysis of (**A**) *S. aureus* (USA300) and (**B**) *S. aureus* (SCO1) treated overnight with 1 mM of D-cystine, reduced with DTT (5 mM), and incubated with designated accessibility probes. Data are represented as mean +/- SD (n = 3).

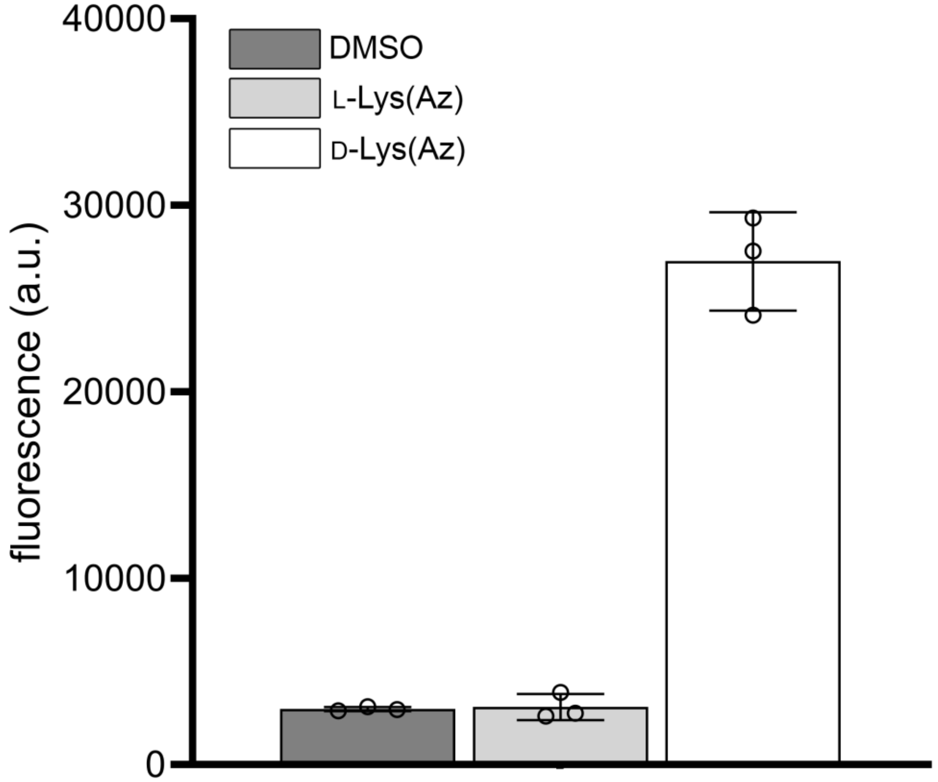


**Figure S5.** Flow cytometry analysis of surface labeled *S. aureus* (ATCC 25923) treated overnight with 1 mM of **D-LysAz** or 1 mM of **L-LysAz** and incubated with 25 μM of **DBCO-Fl**. Data are represented as mean +/- SD (n = 3).

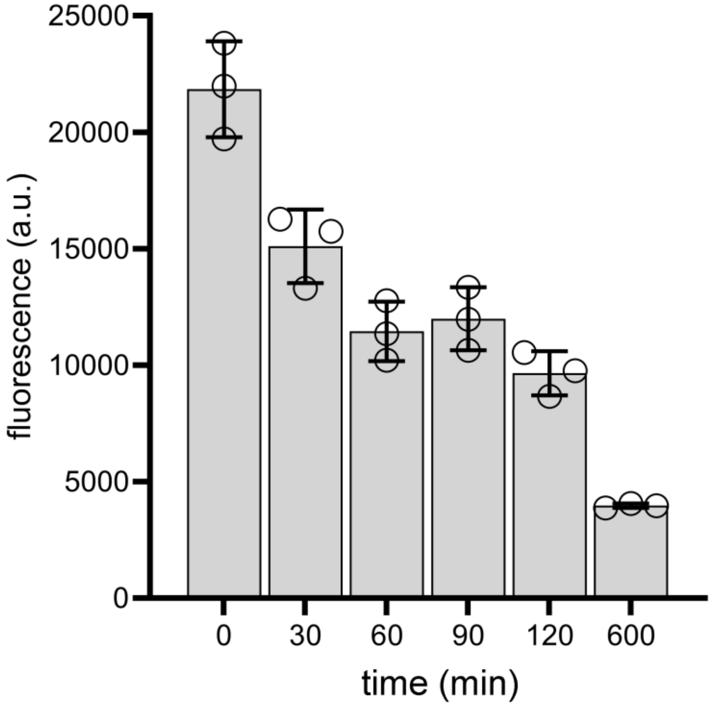


**Figure S6.** Flow cytometry analysis of surface labeled *S. aureus* (ATCC 25923) treated overnight with 1 mM of **D-LysAz** and incubated with 25 μM of **DBCO-Fl**. Cells were fixed with formaldehyde, washed with PBS, and incubated with mutanolysin. Periodically, cells were analyzed by flow cytometry. Data are represented as mean +/- SD (n = 3).

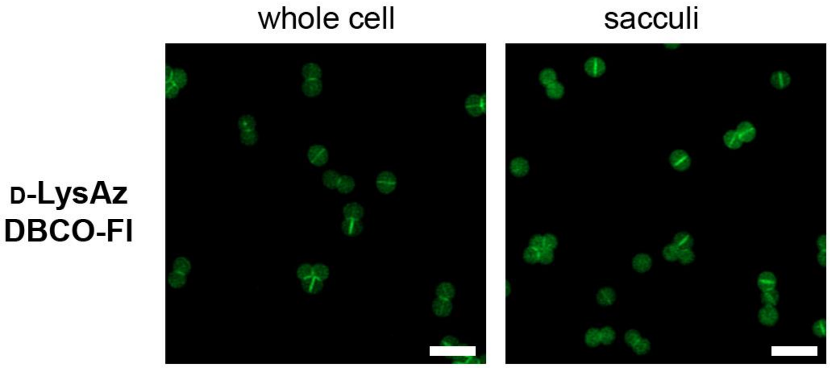


**Figure S7.** Confocal microscopy analysis of surface labeled *S. aureus* (ATCC 25923) treated overnight with 1 mM of **D-LysAz** and incubated with 25 μM of **DBCO-Fl**. Cells were fixed with formaldehyde, washed with PBS, and imaged. Data are represented as mean +/- SD (n = 3).

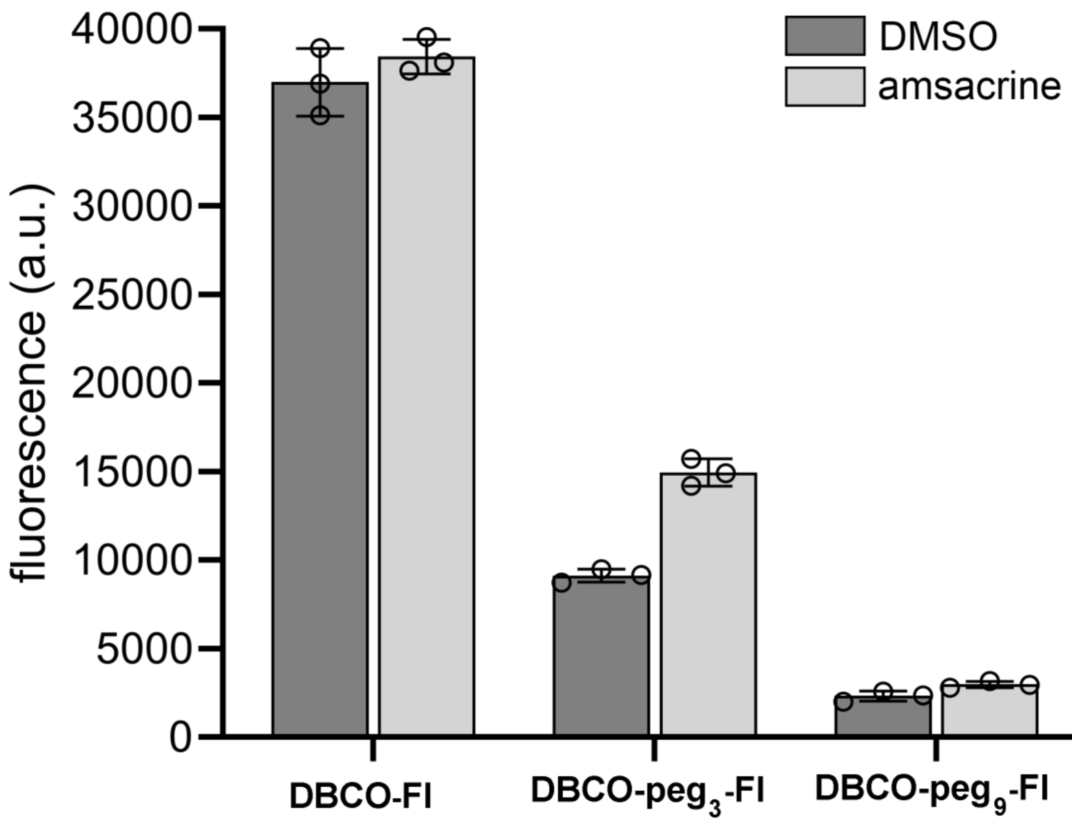


**Figure S8.** Flow cytometry analysis of WT *S. aureus* (ATCC 25923) treated overnight with 1 mM of **D-LysAz,** co-incubated with 0 or 10 μg/mL of amsacrine overnight, followed by a treatment with 25 μM of designated probes.
